# CliPepPI: Scalable prediction of domain-peptide specificity using contrastive learning

**DOI:** 10.64898/2026.03.18.712595

**Authors:** Tanya Hochner-Vilk, Dor Stein, Ora Schueler-Furman, Barak Raveh, Yuh Min Chook, Dina Schneidman-Duhovny

## Abstract

Domain-peptide interactions mediate a significant fraction of cellular protein networks, yet accurately predicting their specificity remains challenging. Peptide motifs typically have short, fuzzy sequence profiles, and their interactions are often weak and transient, limiting the size, coverage, and quality of experimentally validated domain-peptide datasets. Since true non-binders are rarely known, constructing negative examples often introduces bias. While structure-based prediction methods can achieve high accuracy, they are computationally demanding and difficult to scale to the proteome level. We introduce CLIPepPI, a dual-encoder model that leverages contrastive learning to embed domains and peptides into a shared space directly from sequence. Both encoders are initialized from a protein language model (ESM-C) and fine-tuned using lightweight LoRA adapters, enabling parameter-efficient training on positive pairs alone. To overcome data scarcity, we augment ∼3K protein-peptide complexes from PPI3D with ∼150K domain-peptide pairs derived from protein-protein interfaces. CLIPepPI further injects structural information by marking interface residues in the domain sequence, thus guiding the encoders toward binding regions and linking sequence-level learning with structural context. Competitive performance is achieved across three independent benchmarks: domain-peptide complexes from PPI3D, large-scale phage-library data from ProP-PD, and a curated dataset of nuclear export signal (NES) sequences. We demonstrate scalability and generalization through two applications: (i) proteome-wide NES scanning, and (ii) variant-effect prediction, where score changes in domain-peptide interactions between wild-type and mutant sequences discriminate pathogenic from benign variants. Together, CLIPepPI offers a scalable, structure-informed model for predicting domain-peptide specificity and generating meaningful embeddings suited for large-scale proteomic analyses. CLIPepPI is available at: https://bio3d.cs.huji.ac.il/webserver/clipeppi/.

## Introduction

Intrinsically disordered regions (IDRs) are protein segments that lack a stable third-dimensional structure and do not adopt a single defined conformation [1]. This conformational flexibility enables their participation in diverse cellular processes, including signaling and regulation [2, 3]. A major mechanism by which IDRs mediate biomolecular interactions is through short linear motifs (SLiMs). SLiMs are short peptide sequences within IDRs that are recognized by modular binding domains in target proteins [4]. Remarkably, these short peptides mediate approximately 40% of biomolecular interactions [5, 6]. Experimental techniques, such as phage display, have been instrumental in identifying novel binding peptides and characterizing domain-peptide recognition [7–10]. However, these approaches are often time-consuming, costly, and limited in scale, motivating the development of computational methods for predicting domain-peptide interactions.

Predicting domain-peptide specificity is challenging due to subtle sequence features that determine binding specificity, the high sequence variability of SLiMs, and the scarcity of experimentally validated interaction data [11]. SLiMs are highly dynamic, rapidly evolving, and transient binders [12, 13] that typically bind their target domains with low affinity, because only a few residues mediate the interaction [4]. These properties make domain-peptide interactions distinct from general protein-protein interactions, motivating dedicated approaches.

An important class of computational methods for domain-peptide interaction prediction relies on structure-based modeling, such as molecular docking or template-based complex prediction [14–25]. Structure-based interactome prediction pipelines, such as PrePPI [26, 27], extend this idea to proteome-scale inference by matching domains and motifs to structural templates of protein-peptide complexes. With the advent of highly accurate protein structure prediction models [28–31], these approaches can, in principle, be applied directly from sequence data through co-folding of domain with a peptide [32]. For example, fine-tuned protein structure prediction networks have been used to jointly model domain-peptide binding specificity and three-dimensional complex structures [33, 34]. However, the computational cost of structure-based techniques remains prohibitive for proteome-scale screening.

As an alternative, sequence-based prediction methods aim to infer binding specificity directly from the amino acid sequences of the domain and peptide, without requiring explicit 3D structures [35–37]. These approaches leverage representations learned from protein language models or shallow sequence features, enabling rapid and scalable predictions across large datasets. Nevertheless, while sequence-based methods offer efficiency and broad applicability, achieving accuracy comparable to structure-based approaches remains a significant challenge, particularly due to the subtle sequence determinants and the context-dependent nature of domain-peptide recognition. Sequence-based supervised deep learning methods, such as CAMP [38] and HSM [39], have high-accuracy when trained on large datasets of peptides tested using a peptide array designed for a specific domain. The accuracy drops in the general domain-peptide setting, where generalization to previously unseen data is required. The common practice is to use randomly sampled negative domain-peptide pairs for training [38]. However, recent analyses demonstrated that such negative-sampling schemes tend to exploit biases in the input data, rather than learning true biophysical binding specificity [40, 41].

Protein language models (pLMs), trained in an unsupervised manner on millions of protein sequences, learn to predict latent residue embeddings [42–45]. These embeddings encode evolutionary constraints and capture residue-level chemical features, secondary structure properties, and long-range dependencies relevant to protein folding and function. Disobind uses protein language model embeddings to predict partner-dependent IDR interaction interfaces and contacts, outperforming AlphaFold2 and AlphaFold3 on disordered regions [46]. Previous studies have also shown that transfer learning on pre-trained pLMs can be beneficial for specificity prediction [34, 47, 48] especially in settings with limited labeled data. Early work treated pLMs as feature extractors: a frozen encoder followed by a lightweight classifier. These models improved MHC-peptide [34], TCR-peptide, and antibody-antigen [47, 48] specificity prediction. More recently, it was demonstrated that fine-tuning pLMs can boost performance across eight diverse tasks, including interaction prediction and variant effect prediction [49, 50].

The Contrastive Language-Image Pre-training (CLIP) paradigm, originally designed for image-text pairing, trains dual encoders to maximize the cosine similarity of paired inputs while minimizing similarity to all other items within a batch. A key advantage of this approach is that it requires only positive pairs for training, which is beneficial for tasks with limited labeled data. This idea has been adopted for computational biology tasks [51, 52]. For example, PepPrCLIP generates peptide binder candidates and EAGLE designs epitope-specific antibody binders [53, 54]. Additional applications include virtual screening as in DrugCLIP [55] and interaction prediction, as in ImmuneCLIP, ConPLex and FlashPPI [56–59]. These studies suggest that contrastive learning can be adapted for predicting domain-peptide interaction specificity, enabling training on positive interaction data without requiring artificially constructed negative examples typically required in supervised learning.

Building on these advances, we present CLIPepPI, a CLIP-inspired dual-encoder that maps domain and peptide sequences into a shared latent space. Both encoders are initialized from the pLM (ESM-C) and adapted with lightweight LoRA modules, so that only 25% of parameters are updated, keeping training GPU-friendly. We augment our training set using domain-peptide pairs derived from protein-protein interfaces. At inference time, the cosine similarity between the two embeddings is interpreted as a binding score. We evaluate CLIPepPI on three independent test sets and showcase several downstream applications, including variant effect prediction and proteome-wide SLiM scans. We show that the learned embedding space clusters peptides by binding pocket. Together, these results highlight the promise of CLIP-style contrastive learning and parameter-efficient pLM fine-tuning to address long standing data bottlenecks in the prediction of domain-peptide interactions.

## Results

### Overview of CLIPepPI

We developed a data-augmented, sequence-based learning framework that integrates binding site information from known protein-peptide complexes with large-scale augmented data derived from protein-protein complexes. Experimentally validated domain-peptide complexes were collected from PPI3D [60], forming a high-confidence core dataset of 3,000 domain-peptide pairs. To expand the training set, we generated 150,000 augmented domain-peptide pairs derived from protein-protein interaction data in PINDER [61] (Methods). This strategy is motivated by the observation that many globular protein-protein interfaces are dominated by short linear segments that recapitulate most of the native binding energy, making them powerful templates for deriving realistic peptide like interaction patterns [62, 63]. We performed pre-training using the PINDER-derived dataset, and trained using the domain-peptide PPI3D dataset.

Inspired by the CLIP framework [57, 64], we developed CLIPepPI, a contrastive dual-encoder model that predicts binding specificity directly from sequences by independently encoding domains and peptides into a shared structural embedding space (Fig. 1). Both encoders rely on the embedding of ESM-C [65], a state-of-the-art protein language model that captures evolutionary and contextual features of amino acid sequences. To implicitly encode structural information, we augmented the domain sequence with interface residue indicators derived from Voronota analyses [66]. This hybrid representation enables the model to focus on regions most relevant for binding [67, 68] without requiring explicit 3D structures, bridging the gap between structural and sequence-based approaches.

**Fig. 1:**
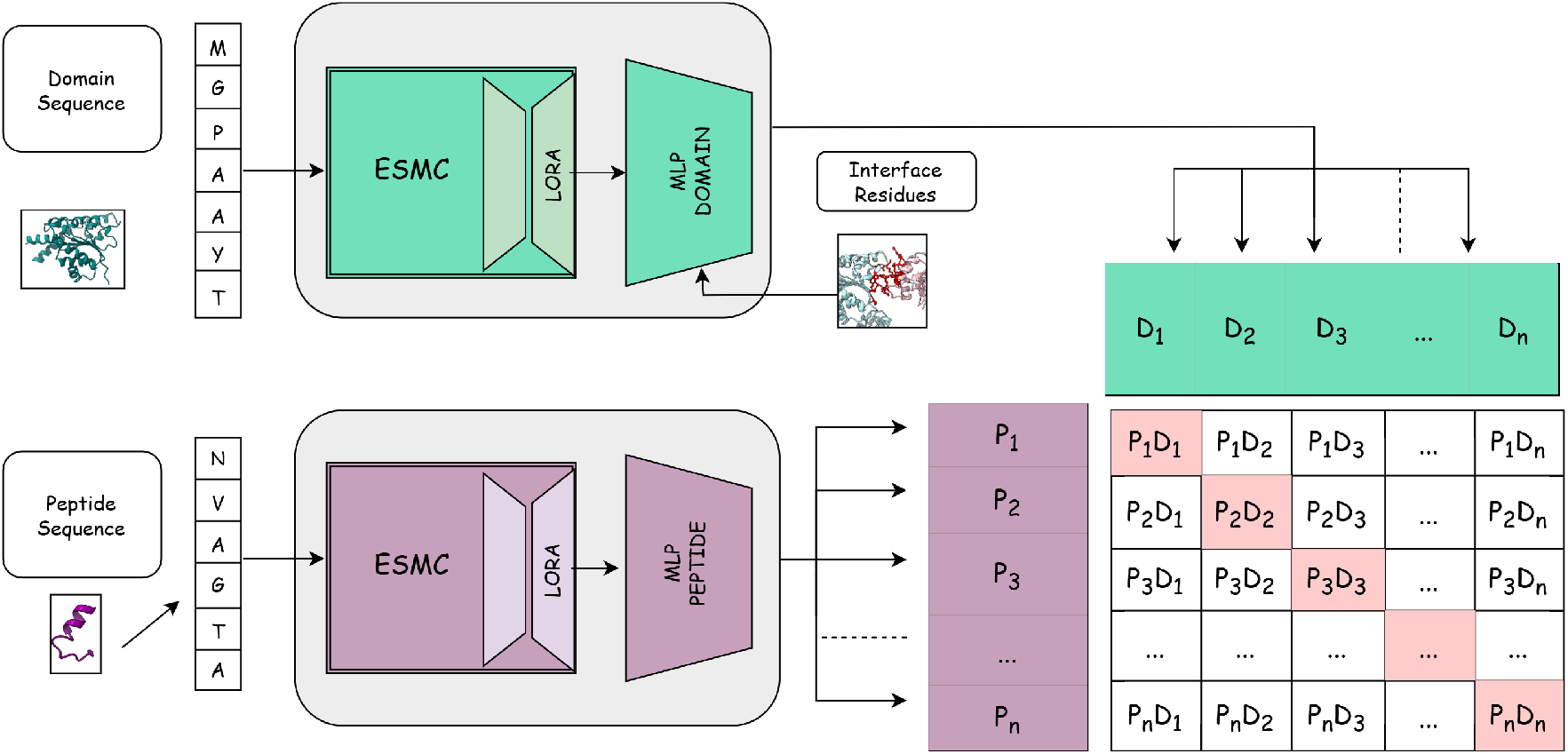
CLIPepPI contrastive-learning framework for domain-peptide recognition. *Top:* Domain sequence is tokenized and passed through a pretrained ESM-C encoder whose weights are frozen except for LoRA adaptation layers (green). A lightweight multilayer perceptron (MLP_domain_) then pools the contextual residue representations and interface indicator into a fixed-length embedding, to yield domain embeddings *D*_1_, &, *D*_*n*_. *Bottom:* In parallel, a peptide sequence is processed by an identical LoRA-adapted ESM-C encoder (purple). The output of ESM-C is passed through the MLP_peptide_ producing a batch of peptide vectors *P*_1_, &, *P*_*n*_. *Right:* For every mini-batch, cosine similarities between all *P*_*i*_ and *D*_*j*_ form an *n × n* score matrix. Cognate domain-peptide pairs (diagonal, red cells) are treated as positives, whereas off-diagonal entries serve as negatives.

Fine-tuning large pretrained protein language models on relatively small domain-peptide datasets risks overfitting and excessive computational cost. To address this, we employed Low-Rank Adaptation (LoRA) [69], which introduces small trainable matrices into the frozen attention layers of the model. LoRA was applied to the query and key projections of the final eight transformer layers, updating approximately 25% of the total parameters. This approach allowed efficient adaptation of ESM-C to the binding prediction task while maintaining parameter efficiency.

Training was performed using a weighted cross-entropy loss, which maximizes cosine similarity between embeddings of true domain-peptide pairs while minimizing it across unrelated pairs within a batch. Importantly, this objective allows training using only positive examples, alleviating the need for unreliable or bias-prone negative labels that are common in protein interaction datasets. Through contrastive learning, the model captures the physicochemical and evolutionary determinants of domain-peptide compatibility. Notably, increasing the contribution of the receptor representation (i.e., using weighted cross-entropy loss) yielded superior performance in datasets where multiple peptides bind the same domain (Methods). In contrast, average cross-entropy loss performed better on our test set which has only a few positive peptides per domain. As real-world applications primarily involve large-scale screening, we therefore adopt the weighted loss formulation for model training.

Once trained, CLIPepPI encodes domains and peptides independently, enabling rapid and scalable binding prediction based on their embeddings similarity. This design allows precomputation of proteome-wide domain embeddings and efficient similarity searches for potential peptide partners. In contrast to structure-based docking or template-based modeling, which require explicit 3D conformations and intensive computation, CLIPepPI offers orders-of-magnitude faster inference while retaining biological interpretability. Consequently, it provides a practical route for large-scale prediction of domain-peptide specificity, proteome-wide SLiM scanning, and variant effect analysis, complementing structure-based methods, such as AlphaFold’s interface-focused confidence metrics, actifpTM [70] or ipSAE [71].

To contextualize our training data and model scope, we adopt the concept of coverage from recent drug-target interaction benchmarking studies [57]. Coverage reflects the average proportion of possible interaction partners for which data exist for each biomolecule. In the context of domain-peptide binding, a low-coverage dataset contains a large and diverse set of domains and peptides, each represented by relatively few annotated interactions, whereas a high-coverage dataset focuses on a narrow subset of biomolecules with many known pairwise interactions. Here, we focus our efforts on a more challenging low-coverage setting to address generalization on previously unseen domains. To achieve this, we train our model on a highly diverse dataset and apply stringent data separation criteria (Methods).

### Specificity prediction

We validate our model using three independent test sets: PPI3D [60], Prop-PD [72–74], and the NES dataset of mutated nuclear export signals [75] (Fig. 2A). After pre-training on the PINDER-derived dataset, we trained seven separate models using the PPI3D dataset. Each model was trained on six out of seven non-overlapping splits (Methods). These seven models were then used to assess generalization performance on the independent PPI3D, ProP-PD and NES datasets, evaluating the robustness of the model across different data splits.

**Fig. 2:**
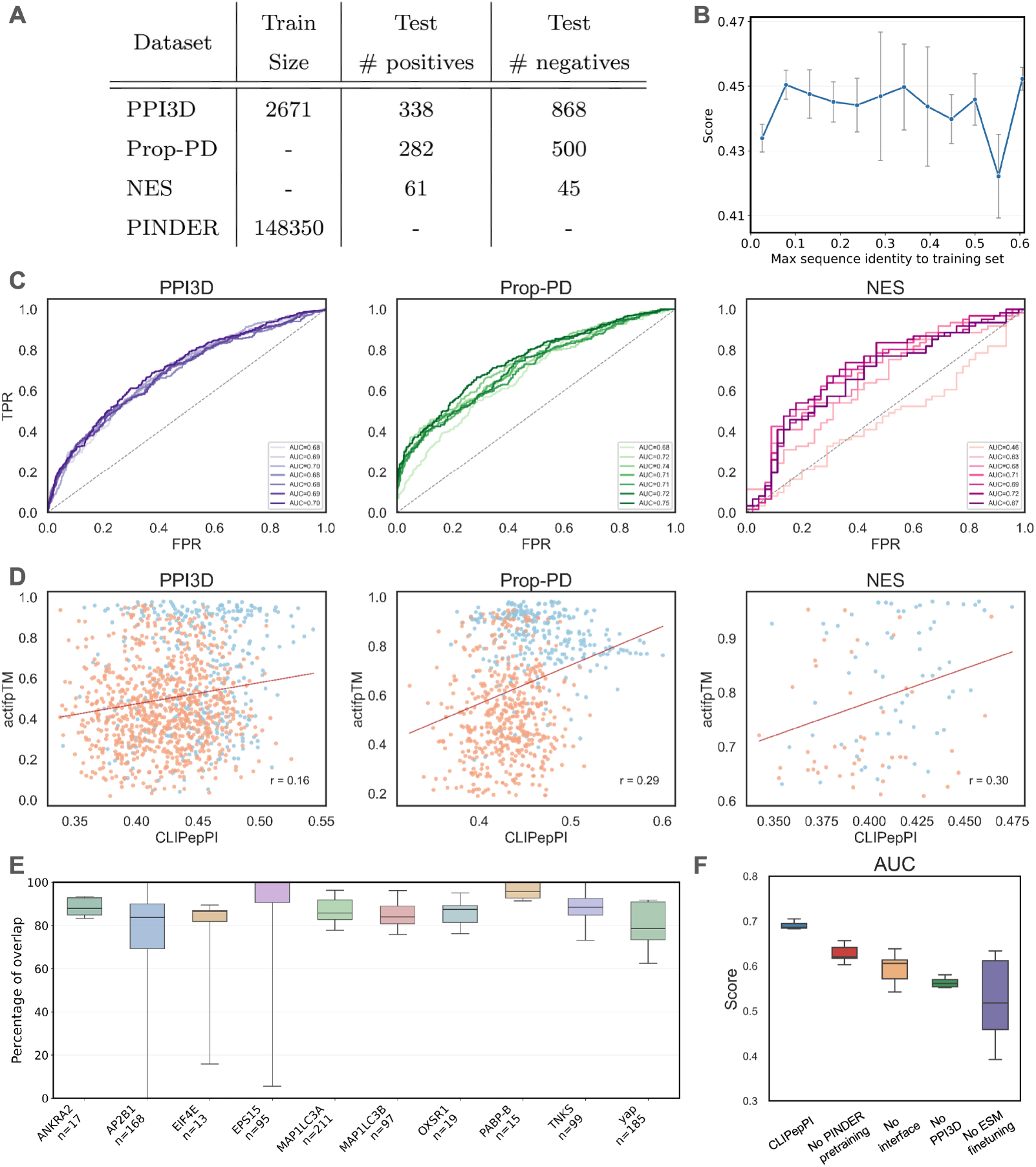
CLIPepPI performance across three benchmark datasets. **A**. Datasets and their sizes. **B**. Model performance (y-axis: cosine similarity scores averaged across folds) vs. test-set similarity to the training set (x-axis: maximum sequence identity of PPI3D test samples to the training set). **C**. Receiver-operating-characteristic (ROC) curves for the seven cross-validated models on the PPI3D, ProP-PD, and NES test sets (left to right). Each shaded curve corresponds to a single cross-validation model; the dashed diagonal denotes random classification. Legends list the per-model area-under-the-curve (AUC) values. **D**. CLIPepPI (x-axis:cosine similarity scores) vs. AlphaFold (y-axis: inter-chain actifpTM confidence scores) for the same datasets. Blue dots represent positive (binding) pairs and orange dots negative pairs. **E**. Percentage of overlap between interface residues identified for the representative peptide and for all other peptides (Prop-PD dataset). Interface residue sets were calculated using Voronota on AlphaFold-predicted structures. **F**. Ablation study for CLIPepPI using the PPI3D test-set. Boxplot AUCs show that removing PINDER pairs, domain interface residue features, PPI3D training data, or ESM fine-tuning with LoRA degrades performance.

#### PPI3D

For non-biased evaluation, we curated a hold-out test set from the PPI3D database containing domain-peptide pairs excluded from all training folds and sharing no sequence identity above the defined thresholds (Methods). Predictions from each of the seven cross-validation models were generated independently and performance metrics were averaged. The model achieved an average AUC of 0.69 ± 0.008 on this dataset (Fig. 2C, Table 1). Performance was largely insensitive to the maximum sequence identity between PPI3D test samples and the training set (Fig. 2B).

**Table 1:** AUC and F1 results across benchmark datasets.

|  | PPI3D |  | ProP-PD |  | NES |  |
| --- | --- | --- | --- | --- | --- | --- |
|  | AUC | F1 | AUC | F1 | AUC | F1 |
| CLIPepPI | $0.69 \pm 0.007$ | $0.45 \pm 0.007$ | $0.72 \pm 0.021$ | $0.54 \pm 0.005$ | $0.65 \pm 0.081$ | $0.71 \pm 0.072$ |
| ipTM | 0.69 | 0.46 | 0.88 | 0.65 | 0.69 | 0.73 |
| actifpTM | 0.68 | 0.45 | 0.91 | 0.58 | 0.74 | 0.73 |
| Combined (ipTM) | $0.72 \pm 0.002$ | $0.47 \pm 0.001$ | $0.90 \pm 0.002$ | $0.65 \pm 0.002$ | $0.71 \pm 0.035$ | $0.73 \pm 0.000$ |
| Combined (actifptm) | $0.72 \pm 0.002$ | $0.45 \pm 0.001$ | $0.93 \pm 0.001$ | $0.58 \pm 0.001$ | $0.75 \pm 0.027$ | $0.73 \pm 0.000$ |

#### ProP-PD

To further evaluate CLIPepPI generalization, we tested it on the ProP-PD database [72– 74] that contains domain-peptide interactions identified by proteomic peptide phage display. This assay detects binding between a specific domain and SLiMs, typically 3-10 amino acids long and located in disordered regions. Such interactions are typically characterized by low-to-mid micromolar affinity and differ from those in the PPI3D dataset, as they are not derived from experimentally determined structures. We included only high-confidence interactions with known binding motifs and annotated domain partners, resulting in a set of 282 pairs from eight domain baits (Fig. 2A). Negative samples for testing were generated as described in Methods. We approximated the domain binding site by predicting the structure of each domain in complex with a representative peptide using AlphaFold2, then identifying interface residues with Voronota [66]. The resulting interface annotation was subsequently applied to all peptides associated with that domain. This simplification is supported by the strong consistency of interface residue sets across peptides for the same domain (Fig. 2E), suggesting that the representative peptide provides a reliable proxy for the shared binding site. Averaged across the seven models, CLIPepPI achieved an AUC of 0.72 ± 0.023 on this dataset (Fig. 2C, Table 1), demonstrating strong generalization despite the increased complexity of the ProP-PD dataset, which captures diverse, weak, and transient interactions.

#### NES evaluation

To assess performance for identification of short nuclear export signal (NES) motifs, we evaluated domain-peptide interactions involving the nuclear export receptor CRM1 and its NES peptides. For this analysis, we used the curated set of human NES-containing proteins reported by Kosugi et al. [75] which contains 106 CRM1-peptide pairs. Unlike the previous datasets, it includes also experimentally validated negatives, providing a robust benchmark for model evaluation. Each of the seven cross-validated checkpoints was applied independently, yielding an average AUC of 0.65 ± 0.084 (Fig. 2C, Table 1). These results illustrate that CLIPepPI remains effective on a small, function-specific dataset.

#### Comparison to AlphaFold

We applied AlphaFold-Multimer [76] and evaluated performance using its confidence metrics for comparison to CLIPepPI scores. The conventional ipTM confidence score is sensitive to the length and disorder of non-interacting flanking segments, masking the real quality of an otherwise identical interface. We therefore computed actifpTM [70, 71], which restricts the calculation to interface residues (and reports per-chain values), providing a length-independent, interaction-focused metric that better reflects the true confidence of the motif-domain contacts central to our domain peptide complexes (Fig. 2D, Table 1). In the PPI3D test set (1,206 pairs), CLIPepPI modestly outperformed actifpTM, with AUC = 0.69 versus 0.68, while the two scores were only weakly correlated (Pearson r = 0.14). In ProP-PD (782 pairs), actifpTM excelled in this set (AUC = 0.90), while CLIPepPI reached 0.72 with a medium positive Pearson correlation (r = 0.32). In the smallest set, NES (106 pairs), actifpTM led with AUC = 0.73 vs. CLIPepPI 0.65 with a medium positive Pearson correlation (r = 0.30). Due to their low correlation, we also evaluated a combined score of the outputs of actifpTM and CLIPepPI, calculating a weighted average: 0.8 · actifpTM + 0.2 · CLIPepPI. The combined scores achieved a slightly higher AUC than either model alone (Table 1). This suggests that the models make complementary predictions, each succeeding on different subsets of examples.

Although the AlphaFold2 actifpTM score outperforms CLIPepPI in some benchmarks, CLIPepPI offers substantial scalability advantages. For example, processing 100 domain-peptide pairs takes approximately 40 minutes with AlphaFold2 on an A40 GPU, whereas CLIPepPI requires only about 1 second. Furthermore, CLIPepPI’s domain and peptide encoders can be directly used to extract embeddings for downstream tasks, making the model broadly useful beyond binding specificity prediction.

#### Ablation experiments

To evaluate the contribution of each component within CLIPepPI, we systematically removed key inputs and training sources (Fig. 2F). Excluding PINDER-derived data (“No PINDER”) leads to a drop in performance, demonstrating the importance of high-quality augmented data. Eliminating interface annotations (“No Interface”) causes further degradation, suggesting that structural interface information contributes valuable context for binding prediction. Excluding the PPI3D evaluation set during training (“No PPI3D”) results in weak performance (AUC *<* 0.6), which is expected since this ablation removes all true domain-peptide training examples, and the test set originates from the same source. Finally, removing ESM fine-tuning with LoRA (“No ESM fine-tuning”) leads to the most dramatic performance decline, emphasizing the crucial role of fine-tuned representations in capturing sequence-level features relevant for domain-peptide specificity.

### Proteome-wide prediction of CRM1-mediated nuclear export signals (NES)

The systematic identification of short linear motifs (SLiMs) that mediate domain-peptide interactions at the proteome scale remains labor-intensive and incomplete, leaving many functional binding sites unannotated. Proteome-wide *in silico* scanning offers a complementary strategy to uncover candidate binding peptides systematically, enabling the generation of testable hypotheses at scale. CLIPepPI is uniquely suited for this task because it operates directly on sequence embeddings, decoupling domain and peptide representation. This allows the domain embedding to be computed once and reused for efficient scoring against millions of peptide segments, making large-scale scanning computationally tractable without sacrificing specificity.

To demonstrate this, we focused on the nuclear export receptor CRM1, which recognizes NES peptides and mediates the nuclear export of numerous proteins. The NES system is particularly suitable for scanning because CRM1-mediated nuclear export is one of the most prevalent peptide-mediated transport mechanisms in human cells. Moreover, numerous NES-containing proteins remain uncharacterized despite their central roles in signal transduction, cell cycle, and regulation [77, 78]. We scanned the entire human reference proteome [79] using a sliding window of 18 amino acids and computed embeddings for every possible segment. Each segment was scored by cosine similarity to the CRM1 domain embedding, and the highest-scoring segments were considered putative NESs and mapped back to their parent proteins.

To validate our scoring scheme, we used a quantitative proteomics study by Kirli et al. [80] classifying ≈ 4, 000 out of 20,0000 human proteins into three categories: (i) cargo proteins (contain NES), (ii) ambiguous, and (iii) non-binders. The top ranked set of proteins (proteome-wide) by CLIPepPI were compared with the experimentally defined CRM1 cargos. Among the top-10 ranking proteins for HeLa cells, five were annotated by Kirli et al. [80] as experimentally validated or ambiguous CRM1 cargoes (three and two proteins, respectively) (Fig. 3). Even among the top-4000 proteins, there were very few annotated non-binders. Similar results were obtained with the annotated dataset of *Xenopus laevis* oocytes. These findings highlight the potential utility of our approach for proteome-wide discovery of binding peptides.

**Fig. 3:**
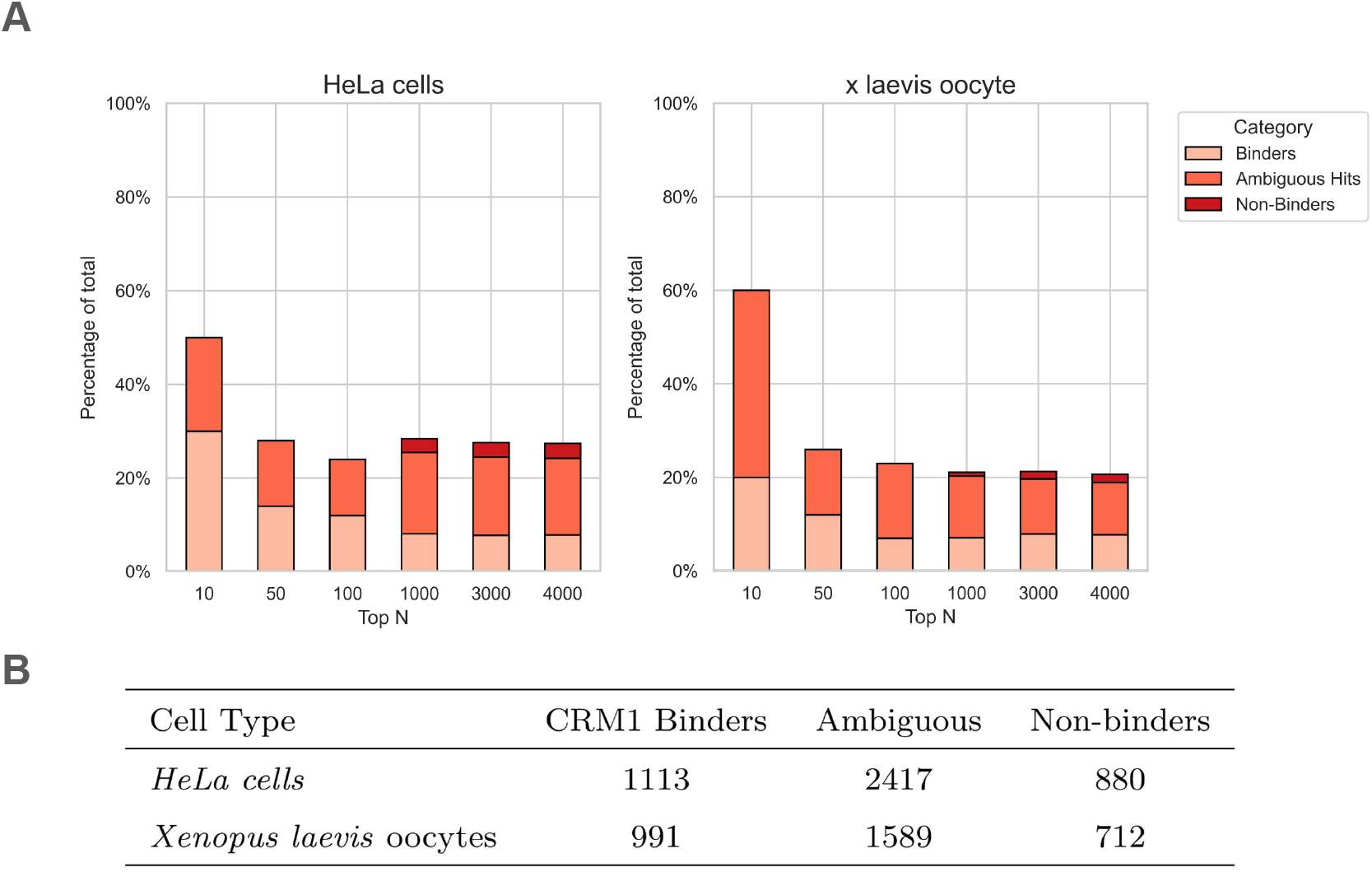
Proteome-wide scan for NES motif containing proteins compared against experimental proteomics annotations of *HeLa* cells and *Xenopus laevis* oocytes [80]. **A**. For increasing Top-*N* cutoffs, stacked bars report the percentage of total hits among the highest-scoring candidates, partitioned by annotation categories from proteomics experiments (see Legend). **B**. Summary of results for proteomics experiments by Kirli et al. [80].

### Pathogenicity analysis

Another important application of CLIPepPI is the mechanistic interpretation of variant effects on protein interactions. To explore this, we analyzed missense mutations from ClinVar [81] that map to three domain families (SH2, SH3 and PDZ) and general domains from our held-out test set (PPI3D), compiled by F. Brunello et al. [82]. Binding peptides for these domains were obtained from the Protein Binding Domain (PBD) dataset [39] and PPI3D [60], resulting in 128 classified domain variants and 3,592 peptides with 7,525 domain-peptide pairs. For each variant, we computed the difference in CLIPepPI scores between the wildtype and mutant domain. Because CLIPepPI explicitly models domain-peptide interaction compatibility, differences in score can be interpreted as changes in binding propensity. This enables mechanistic discrimination between variants that primarily perturb interaction interfaces and those whose pathogenic effects are likely due to other mechanisms, such as altered folding stability. Consistent with previous observations that disease-associated mutations are enriched at interaction interfaces [83, 84], we observed significantly larger score changes for pathogenic compared to benign variants, across all four domain families (Fig. 4). Importantly, CLIPepPI is not a general variant-effect or pathogenicity predictor, as pathogenicity can arise from mechanisms beyond altered recognition. Instead, it provides an interaction-specific, mechanistically interpretable signal that helps distinguish variants likely to perturb domain-peptide binding from those acting through other effects.

**Fig. 4:**
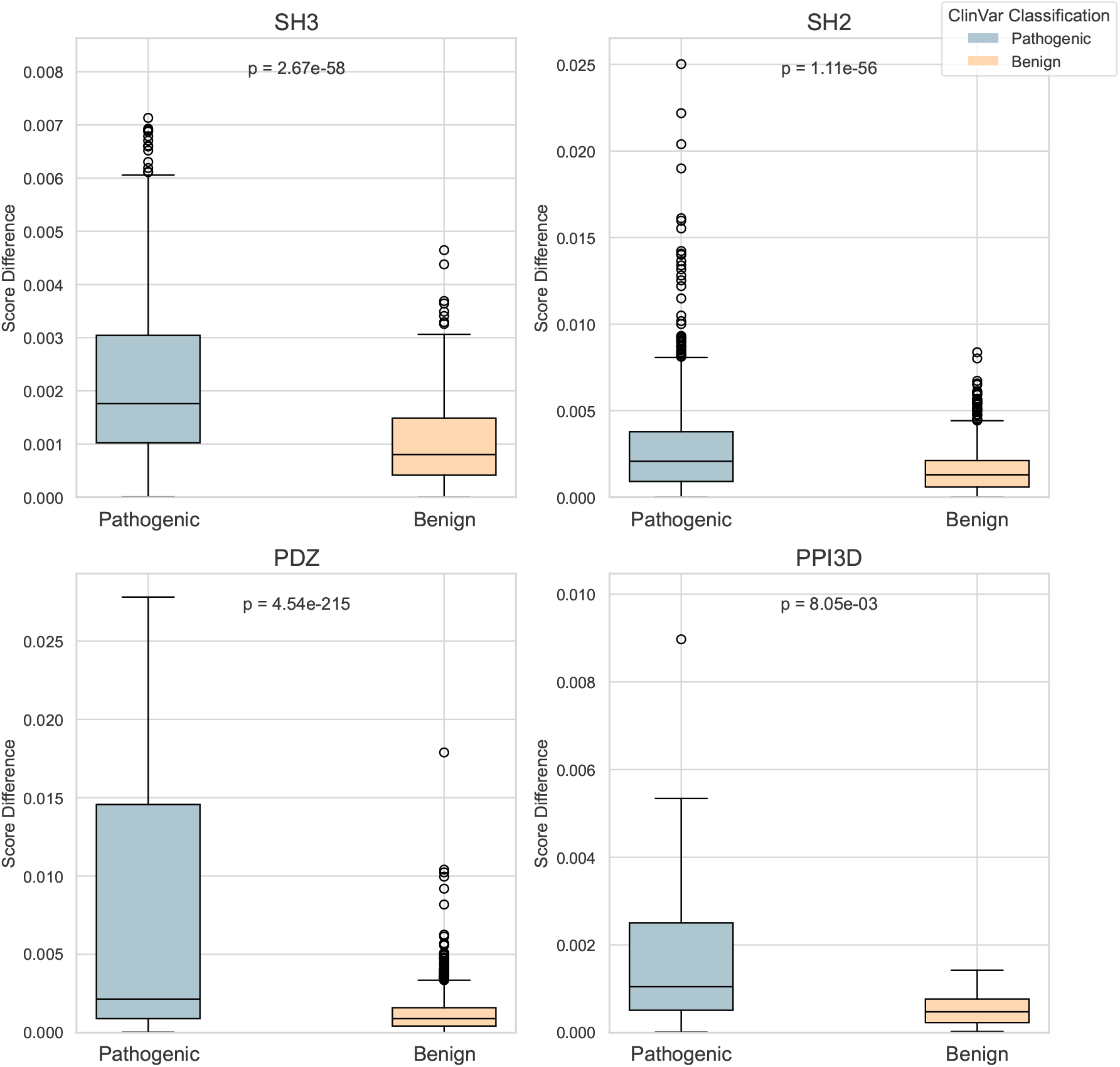
Analysis of effects of ClinVar variants. Distributions of score difference (wildtype vs. mutant) for three domain families (SH3, WW, PDZ) and general domains (PPI3D). Boxplots show the median, interquartile range; points are outliers. T-test *p*-values are annotated above each panel, indicating a significant separation between the score differences of pathogenic and benign variants.

### 2D visualization of peptide embedding

To further evaluate the learned peptide embedding space, we performed a two-dimensional t-SNE analysis of peptide embeddings for the Prop-PD and PBD databases [39, 72] (Fig. 5). The projection reveals distinct clusters that correspond to known binding preferences. For example, in the PBD, which is a domain database, the PDZ-binding peptides form a tight, isolated cluster, while SH2 and SH3 have broader overlaps with other domains. These maps demonstrate that the representation space encodes domain-specific binding features.

**Fig. 5:**
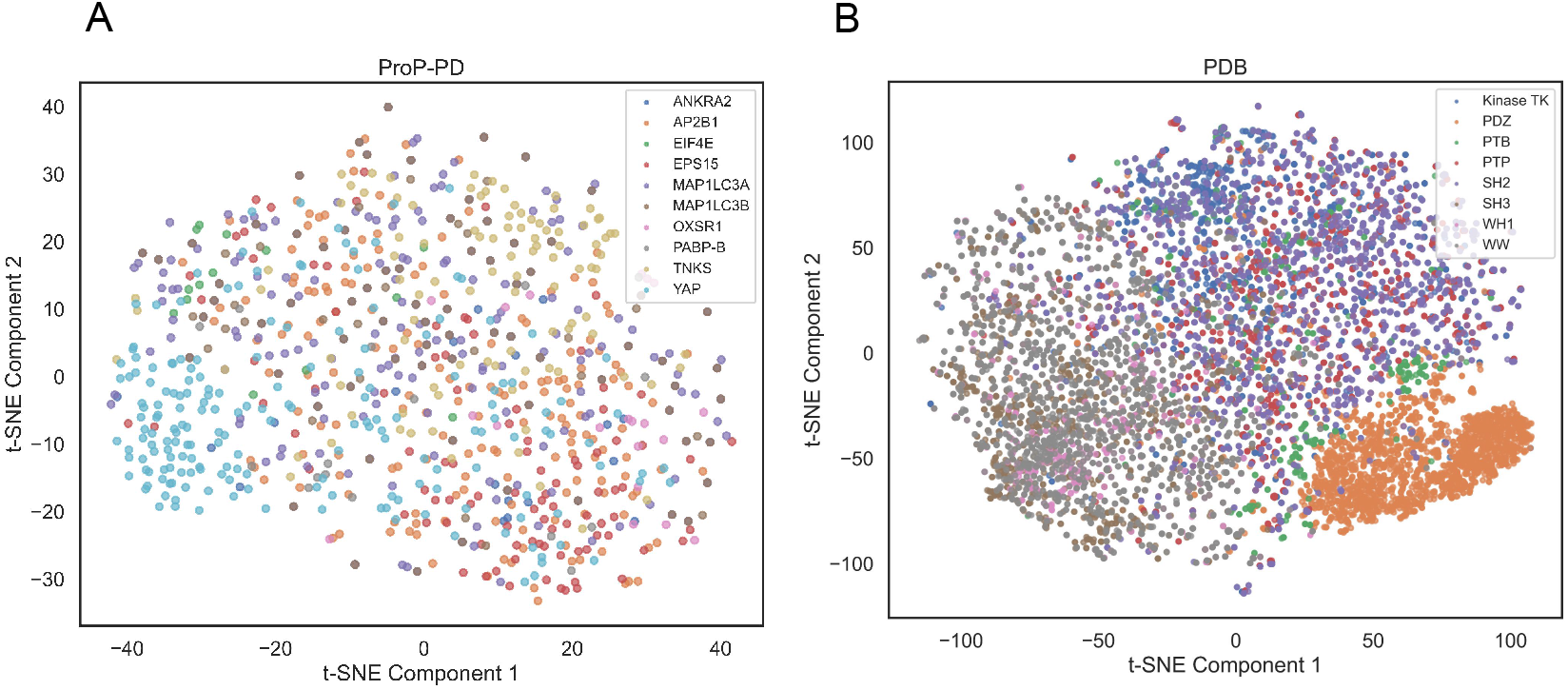
t-SNE projections of domain-specific peptide embeddings. Each point represents a single peptide encoded by the CLIPepPI model and embedded in 900-dimensional space; points are colored by binding domain shown in the legends in each panel. **A**. Peptides from the ProP-PD screen cluster in part according to their cognate receptors (e.g., ANKRA2, AP2B1, EIF4E), indicating that the learned representation captures receptor-specific binding preferences. **B**. Peptides from the PBD database cluster by receptor-domain family (PDZ, SH3, SH2, PTB, PTP, WH1, WW, and Kinase TK), with compact clusters for highly selective domains such as PDZ, SH2 and SH3, and broader overlaps for more promiscuous classes like WW domains. Axes show the first and second t-SNE components. No additional supervision was applied during projection, demonstrating that specificity information is inherently encoded in the learned embedding space.

### Comparative CLIPepPI performance across PBD families

Drug-target interaction studies have highlighted that the coverage of interaction data, defined as the proportion of possible partners with known interactions, strongly shapes the type of modeling problem to be solved [57]. In low-coverage regimes, each biomolecule is sparsely annotated, and models must generalize broadly across diverse families, including previously unseen families. In contrast, high-coverage regimes contain dense blocks of interactions within a limited biochemical space, enabling fine-grained discrimination between closely related molecules. These correspond to two distinct but complementary applications: proteome-scale scanning versus optimization within a specific biochemical context.

We have constructed our training set to enable CLIPepPI to work in a more challenging low-coverage regime. Specifically, we have used non-redundant PINDER and PPI3D datasets (Methods). To assess CLIPepPI’s applicability to a high-coverage subproblem, where specialized models can leverage richer local features to enhance specificity, we fine-tuned separate versions of the model for each domain family, using the same training set as in the HSM/D study [39].

To evaluate CLIPepPI in this regime, we compared its finetuned models performance to HSM/D [39] and CAMP [38], both supervised models, across eight major PBD families (Fig. S1). With the exception of the Kinase TK family, AUC results ranged between 0.84 and 0.96. CLIPepPI outperformed CAMP on SH3 and PTP domains, surpassed HSM on SH2 domains, and achieved slightly lower scores on the remaining families. Overall, these results highlight CLIPepPI’s ability to deliver competitive predictions across diverse peptide-binding domain families in a high-coverage regime.

## Discussion

The CLIPepPI model introduced in this study leverages contrastive learning and fine-tuned pLMs to predict domain-peptide interactions (Fig. 1). Using only positive samples and dual encoders inspired by the CLIP architecture, CLIPepPI efficiently maps peptide and domain sequences into a shared embedding space. Through evaluation of different benchmark data sets, PPI3D, ProP-PD, and NES, we demonstrate that CLIPepPI effectively captures specificity in various biological contexts. The embeddings generated by our model capture informative representations (Fig. 5) and enable diverse downstream applications. Here, we demonstrate their utility for mutation variant analysis (Fig. 4) and proteome-wide scanning for binding motifs (Fig. 3).

Our approach uniquely combines contrastive learning with structural data for peptide-protein binding prediction, similar to the pocket encoder in DrugCLIP [55]. By augmenting domain sequences with interface residue annotations derived from structural data, the model captures spatial binding context without requiring explicit 3D modeling at inference time. This integration of sequence-based representation learning and structure-informed supervision bridges the gap between sequence- and structure-based models.

Equally critical to CLIPepPI’s success is its fine-tuning strategy. The model employs LoRA to finetune a pretrained state-of-the-art ESM-C pLM, allowing only a fraction of the parameters to update. This approach preserves the rich biochemical priors learned from massive unlabeled sequence datasets while adapting the model to the nuances of domain-peptide recognition. Fine-tuning proved essential for capturing subtle physicochemical features underlying binding specificity. Indeed, ablation experiments revealed that removal of ESM-C fine-tuning caused the most substantial drop in performance (Fig. 2F). Since PLMs are trained on single protein sequences, finetuning is necessary to learn interaction representations. This result emphasizes how contrastive learning on fine-tuned embeddings enables the model to learn generalizable, high-resolution representations of molecular interactions that enable prediction in a low-coverage setting.

One of the main challenges encountered in the development of CLIPepPI was the inherent data scarcity in experimentally validated domain-peptide pairs and structural data, which limited the quality and diversity of the training data. The data augmentation strategy, combining high-confidence structural complexes from PPI3D with pairs derived from PINDER protein-protein interfaces, was instrumental in overcoming the scarcity of experimentally validated domain-peptide data. This hybrid training dataset increased diversity while preserving physical realism, improving generalization to unseen domains and peptides. Future extensions could incorporate additional augmented examples by extracting peptide-like fragments from domain-domain interaction interfaces or by segmenting monomeric proteins into putative domain-peptide pairs through cleavage of N- or C-terminal regions [20]. Additionally, data can also be obtained from high-confidence AlphaFold predictions.

A key advantage of CLIPepPI is its computational scalability for proteome-wide analysis. Although CLIPepPI shows strong predictive performance, AlphaFold’s actifpTM metric occasionally outperforms it. However, CliPepPI is orders of magnitude faster than structure-based methods like AlphaFold for interaction prediction. This efficiency enables practical applications at a proteomic scale, such as scanning the entire human proteome for NES peptides (Fig. 3), demonstrating how embedding-based scanning can accelerate large-scale discovery of short linear motifs that mediate transient, low-affinity interactions.

Beyond motif discovery, CLIPepPI provides a quantitative framework for the interpretation of variant effects. Differences in embedding-space similarity between wild-type and mutant sequences correlated strongly with known pathogenicity across multiple domain families, indicating that the model captures sequence perturbations that disrupt molecular recognition (Fig. 4). This capability highlights the potential of contrastive embeddings as a general tool for interpreting the functional consequences of mutations in signaling and regulatory proteins.

In conclusion, CLIPepPI presents an innovative and computationally efficient solution for predicting the specificity of domain-peptide interactions, effectively overcoming traditional data bottlenecks by harnessing contrastive learning with positive samples only and lightweight fine-tuning strategies. The ability of CLIPepPI to rapidly provide meaningful biological predictions positions it as a promising tool for large-scale biological analyses and facilitates deeper insights into peptide-mediated cellular processes.

## Methods

### Datasets

The training data were derived from protein structures originating from two main sources. The first source was the PPI3D database [60], which curates experimentally determined protein-peptide complexes from the Protein Data Bank (PDB) [85]. These pairs represent biologically validated interactions and were used directly for training and evaluation. However, the number of available protein-peptide structures in the PDB is relatively small (*<* 4,000), limiting its utility as a standalone training resource.

To address this data gap, we augmented the dataset with protein-peptide pairs derived from the PINDER database of protein-protein interactions (PPIs) [61]. A total of 150,000 pairs were randomly sampled uniformly across all clusters annotated in PINDER to maximize diversity within the augmented dataset. During data generation (Fig. 6), each complex was first downloaded from PINDER, annotated such that the larger chain served as the receptor and the smaller as the ligand. Interface residues were identified using Voronota, a geometric tool to detect atomic contacts based on Voronoi diagrams [66]. We then extracted a short peptide fragment from the ligand chain by scanning all continuous 9-14 residue subsequences that intersected the interface and choosing the subsequence with the highest interface-residue fraction. Complexes were discarded if no subsequence contained at least 60% interface residues. The resulting augmented protein-peptide complexes, combined with the true complexes from PPI3D, provided a richer dataset for model training and testing.

**Fig. 6:**
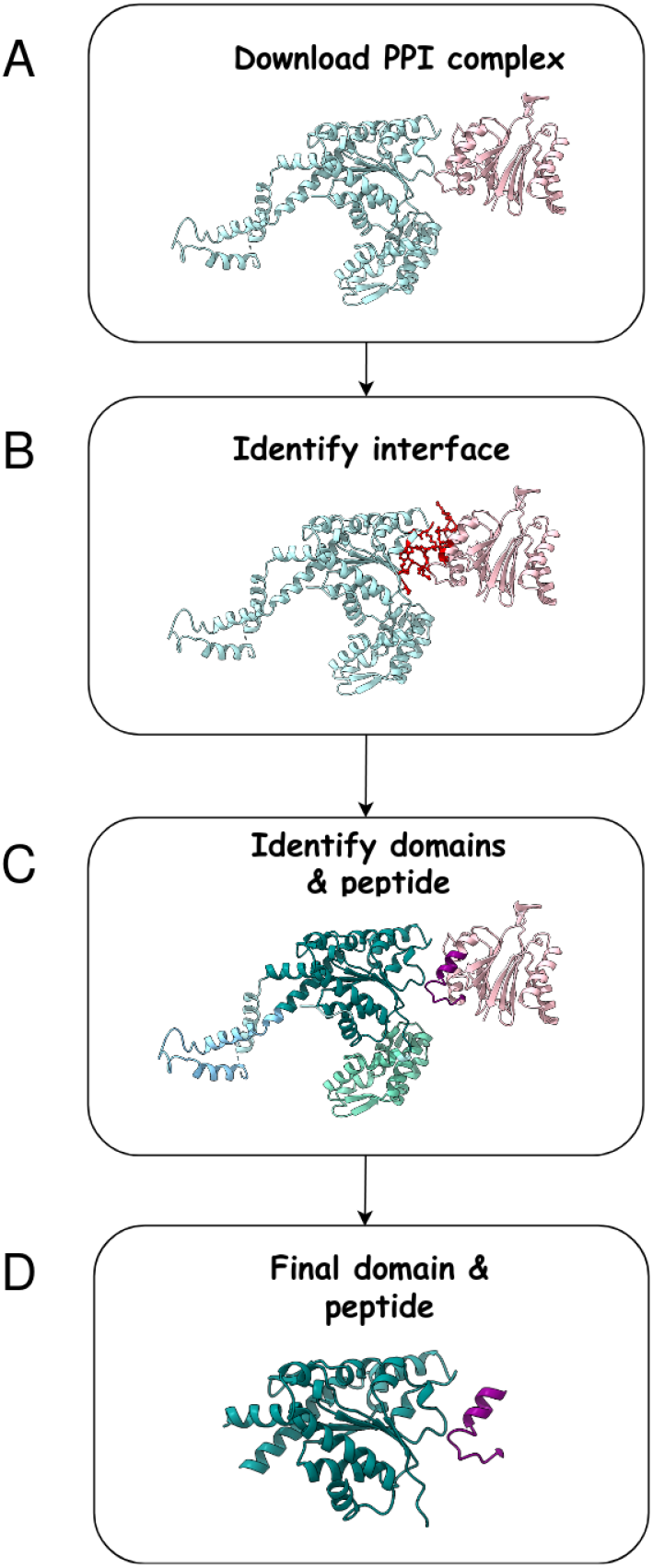
Pipeline for extracting domain-peptide pairs from protein-protein complexes. **A**. A complex is first downloaded from PINDER and the larger chain is defined as a receptor while the smaller as a ligand. **B**. The ligand interface contacts (red) are detected by Voronota. **C**. The receptor chain is segmented into subdomains using Chainsaw. A continuous stretch of 9-14 residues that covers the majority of the interface is extracted as the peptide ligand, and the receptor is truncated to the single domain that participates in the interface. Distinct colors mark individual chains or domains. **D**. The final receptor domain (cyan) and ligand peptide (magenta) whose sequences are provided to the CLIPepPI training pipeline.

To restrict the input to the most relevant region, receptors were represented by their binding domains rather than whole protein chains. Focusing on the domain reduced sequence length, ensured compatibility with the model’s fixed input size constraints, and emphasized the structural region interacting with the peptide. This representation also improved computational efficiency by reducing runtime. The binding domains were trimmed to include up to 400 amino acids in length. Samples of less than 400 amino acids were padded. This trimming was performed using Chainsaw, a deep learning-based method to predict the boundaries of the structural domain [86]. Using the parsed domains, we chose all the Chainsaw predicted subdomains that have residues in the interface. This was particularly beneficial for cases where the binding region was discontinuous in sequence. The maximum length for a peptide was set to 28.

The final dataset contains 3,231 natural complexes from PPI3D and 150,000 derived protein-peptide binding pairs from PINDER. The PPI3D data set was divided into train and test sets using a 9:1 ratio, with sequence identity cutoff of 60% for the domain and 75% for the peptide to prevent data leakage. We used the same cutoffs to generate the 7 non-overlapping splits for training the models.

### Structure prediction with AlphaFold

AlphaFold-Multimer v.2.3 [76] runs were performed using ColabFold [87] with default parameters. The model’s top predicted structure was used for ipTM- and actifpTM-based specificity prediction (Fig. 2D), as well as for the evaluation of the generality of interface residue selection (Fig. 2E).

### Encoder architecture

Inspired by the OpenAI CLIP framework [64], we used a dual encoder contrastive learning approach. CLIP jointly trains an image encoder and a text encoder so that matched image-caption pairs are projected close together, and mismatched pairs far apart, all in a shared embedding space. This approach is a self-supervised method and requires only positive pairs for training data, a great advantage in our case. Analogously, we train two independent encoders: one for peptide sequences and one for domain sequences. Each encoder produces an embedding in a shared latent space of fixed dimensionality. The encoders were trained to maximize the similarity between embeddings of true domain-peptide pairs, while minimizing the similarity between non-interacting domain-peptide pairs.

Both domain and peptide encoders are based on the ESM-C model of evolutionary scale modeling [65], a pLM built on a Transformer architecture. Evolutionary Scale reports that ESM-C is designed as a drop-in replacement for ESM2 [43] and comes with major performance benefits of 300M parameters vs. ESM2’s 650M parameters and delivers similar performance.

Each encoder is followed by a small multilayer perceptron (MLP) head that produces a fixed-length embedding. The MLP receives as input the sequence embedding produced by ESM-C. This embedding is processed through two feedforward layers with ReLU activations, yielding a final sequence representation of dimension 900. The domain and peptide encoders share this architecture, but operate independently and do not share weights. For the domain encoder, we additionally provide a one-hot vector indicating which residues participate in the interface, as identified by Voronota [66]. This feature adds structural information to the domain embedding and is beneficial for binding prediction as demonstrated in our ablation analysis (Fig. 2F).

### Loss function

Each training batch consists of *N* domain peptide pairs. The model outputs *N* domain embeddings and *N* peptide embeddings, each in the shared latent space. A pairwise cosine similarity matrix *S* ∈ *R*^*N*×*N*^ is computed between all domain and peptide embeddings (Eq. 1). In this matrix, the diagonal entries correspond to the true pairs of domain peptides, while the off-diagonal entries represent negative pairs (non-interacting) (Fig. 1).

To train the model, we used a weighted cross-entropy loss. Specifically, each row of the similarity matrix is treated as a softmax distribution over peptides for a given domain and vice versa for each column. The final loss is a weighted average of the cross-entropy for both peptide and domain (Eq. 4). Emphasizing the contribution of the receptor by introducing weighting between domain-to-peptide and peptide-to-domain losses improved screening performance (Figs. S2-S5).

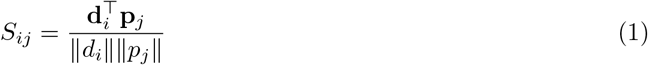

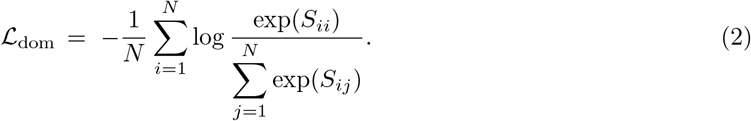

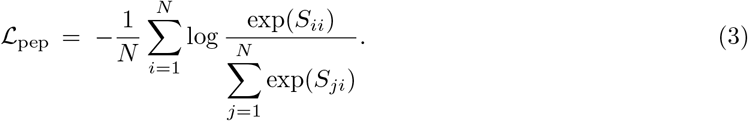

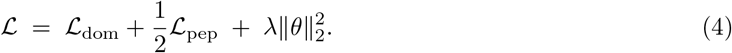

### Fine-tuning with LoRA

We train both the ESM-C parameters and the MLP heads to enable the model to better capture features relevant to our specific task. Finetuning the ESM-C encoder significantly improved binding prediction performance, as shown in the ablation results (Fig. 2F). However, given the large number of parameters in ESM-C, full fine-tuning is computationally expensive and memory intensive. To address this, we adopt LoRA (Low-Rank Adaptation), a parameter-efficient fine-tuning method that introduces trainable low-rank matrices into the attention layers of a frozen pretrained model [69]. Our model was fine-tuned using LoRA adapters during training for the domain-peptide binding task. LoRA adapters were inserted into the last eight layers of ESM-C, allowing 25% of the total model parameters to be trained while keeping the rest of the network frozen. This technique enables adaptation to new tasks by updating only a small subset of parameters, significantly reducing memory and compute requirements. In our setup, LoRA was applied to the query and key attention projections with a rank of 8 and scaling factor (alpha) of 32. By restricting the number of trainable weights, LoRA also helps mitigate the risk of overfitting, especially when working with limited labeled data.

To further improve generalization, we combined weight decay (L2 regularization) with dropout during training. We also improved robustness by randomly masking 10% of the amino acid positions in every input sequence: selected residues were replaced with the ‘-’ token, which the ESM-C encoder interprets as an unknown residue. Masking those residues serves as input-level regularization: by hiding 10% of the amino acid identities, we inject noise that prevents the model from simply memorizing exact sequence patterns seen during training. Instead, the encoder must rely on the surrounding context to construct a useful representation, which typically leads to better generalization on unseen sequences.

### Negative sample generation

For validation and initial testing, we created negative samples, by randomly pairing peptides with domains that do not form known interactions. A negative pair was included only if (i) the domain shared less than 50% sequence identity with the original (true) domain and (ii) the peptide shared less than 80% identity with all peptides that bind to that domain. Furthermore, both the domain and the peptide in each negative pair were excluded from the training set to ensure the evaluation on previously unseen sequences.

### ClinVar data

To evaluate the utility of CLIPepPI for variant effect interpretation, we compiled a dataset of missense mutations from ClinVar, a publicly curated repository that aggregates clinically annotated genetic variants [81]. We used the preprocessed ClinVar dataset provided by F. Brunello et al. [82] and extracted three domain families with annotated mutations: SH2, SH3 and PDZ. We also included general domains from our PPI3D test set. For each domain sequence, we paired the wild-type domain with its experimentally validated binding peptides obtained from the PBD database [39] or PPI3D [60], correspondingly. This procedure yielded 128 classified variants and 3,592 peptides, forming 7,325 domain-peptide pairs.

## Code and data availability

CLIPepPI is implemented using PyTorch. The code and model weights are available as part of the repository https://github.com/dina-lab3D/CLIPepPI. The datasets that were used in training and evaluating CLIPepPI are available at https://huggingface.co/datasets/dorstein0909/ClipPepPI/tree/main. CLIPepPI webserver is available at: https://bio3d.cs.huji.ac.il/webserver/clipeppi/.

## Acknowledgments

We are grateful for the support of the Center for Interdisciplinary Data Science Research (CIDR) at the Hebrew University of Jerusalem. In particular, we would like to thank Haimasree Bhattacharya from CIDR for her contributions to web server of CLIPepPI. O.S-F is supported by the Israel Science Foundation (ISF 301/2021) founded by the Israel Academy of Sciences and Humanities (3091/2021). B.R. is supported by an Israel Science Foundation (ISF 385/24). D.S and T. H. are supported by the Israeli Science Foundation (ISF 737/2023) and the European Research Council (ERC) under the European Union’s Horizon Europe research and innovation programme (ERC Consolidator Grant 101171306). Molecular graphics and analyses performed with UCSF ChimeraX, developed by the Resource for Biocomputing, Visualization, and Informatics at the University of California, San Francisco, with support from National Institutes of Health R01-GM129325 and the Office of Cyber Infrastructure and Computational Biology, National Institute of Allergy and Infectious Diseases.

## Notes

### Competing Interest Statement

The authors have declared no competing interest.

### Summary of Updates

We have updated the TK dataset for PBD using the 2023 version. As a result Figures S4 and S5 were updated.

## References

[1] Tompa, P.: Intrinsically unstructured proteins. Trends in biochemical sciences 27(10), 527–533 (2002)

[2] Dyson, H.J., Wright, P.E.: Intrinsically unstructured proteins and their functions. Nature reviews Molecular cell biology 6(3), 197–208 (2005)

[3] London, N., Movshovitz-Attias, D., Schueler-Furman, O.: The structural basis of peptide-protein binding strategies. Structure 18(2), 188–199 (2010)

[4] Tompa, P., Davey, N.E., Gibson, T.J., Babu, M.M.: A million peptide motifs for the molecular biologist. Molecular cell 55(2), 161–169 (2014)

[5] Neduva, V., Linding, R., Su-Angrand, I., Stark, A., Masi, F.d., Gibson, T.J., Lewis, J., Serrano, L., Russell, R.B.: Systematic discovery of new recognition peptides mediating protein interaction networks. PLoS biology 3(12), 405 (2005)

[6] Mondal, A., Chang, L., Perez, A.: Modelling peptide–protein complexes: docking, simulations and machine learning. QRB discovery 3, 17 (2022)

[7] Sidhu, S.S., Lowman, H.B., Cunningham, B., Wells, J.A.: Phage display for selection of novel binding peptides. Methods in enzymology 328, 333–63 (2000)

[8] Davey, N.E., Seo, M., Yadav, V.K., Jeon, J., Nim, S., Krystkowiak, I., Blikstad, C., Dong, D., Markova, N., Kim, P.M., Ivarsson, Y.: Discovery of short linear motif-mediated interactions through phage display of intrinsically disordered regions of the human proteome. The FEBS Journal 284 (2017)

[9] Gogl, G., Jane, P., Caillet-Saguy, C., Kostmann, C., Bich, G., Cousido-Siah, A., Nyitray, L., Vincentelli, R., Wolff, N., Nomine, Y., et al.: Dual specificity PDZ-and 14-3-3-binding motifs: a structural and interactomics study. Structure 28(7), 747–759 (2020)

[10] Garai, Á., Zeke, A., Gógl, G., Törő, I., Fördős, F., Blankenburg, H., Bárkai, T., Varga, J., Alexa, A., Emig, D., et al.: Specificity of linear motifs that bind to a common mitogen-activated protein kinase docking groove. Science signaling 5(245), 74–74 (2012)

[11] Van Roey, K., Uyar, B., Weatheritt, R.J., Dinkel, H., Seiler, M., Budd, A., Gibson, T.J., Davey, N.E.: Short linear motifs: ubiquitous and functionally diverse protein interaction modules directing cell regulation. Chemical reviews 114(13), 6733–6778 (2014)

[12] Davey, N.E., Van Roey, K., Weatheritt, R.J., Toedt, G., Uyar, B., Altenberg, B., Budd, A., Diella, F., Dinkel, H., Gibson, T.J.: Attributes of short linear motifs. Molecular BioSystems 8(1), 268–281 (2012)

[13] Raveh, B., Karp, J.M., Sparks, S., Dutta, K., Rout, M.P., Sali, A., Cowburn, D.: Slide-and-exchange mechanism for rapid and selective transport through the nuclear pore complex. Proceedings of the National Academy of Sciences 113(18), 2489–2497 (2016). 10.1073/pnas.1522663113

[14] London, N., Raveh, B., Cohen, E., Fathi, G., Schueler-Furman, O.: Rosetta flexpepdock web server—high resolution modeling of peptide–protein interactions. Nucleic acids research 39(suppl 2), 249–253 (2011)

[15] Charitou, V., Van Keulen, S.C., Bonvin, A.M.: Cyclization and docking protocol for cyclic peptide– protein modeling using haddock2.4. Journal of chemical theory and computation 18(6), 4027–4040 (2022)

[16] Raveh, B., London, N., Schueler-Furman, O.: Sub-angstrom modeling of complexes between flexible peptides and globular proteins. Proteins: Structure, Function, and Bioinformatics 78(9), 2029–2040 (2010)

[17] Raveh, B., London, N., Zimmerman, L., Schueler-Furman, O.: Rosetta FlexPepDock ab-initio: simultaneous folding, docking and refinement of peptides onto their receptors. PloS one 6(4), 18934 (2011)

[18] Smith, C.A., Kortemme, T.: Structure-based prediction of the peptide sequence space recognized by natural and synthetic PDZ domains. Journal of molecular biology 402(2), 460–474 (2010). 10.1016/j.jmb.2010.07.032

[19] Alam, N., Goldstein, O., Xia, B., Porter, K.A., Kozakov, D., Schueler-Furman, O.: High-resolution global peptide-protein docking using fragments-based PIPER-FlexPepDock. PLoS computational biology 13(12), 1005905 (2017)

[20] Khramushin, A., Ben-Aharon, Z., Tsaban, T., Varga, J.K., Avraham, O., Schueler-Furman, O.: Matching protein surface structural patches for high-resolution blind peptide docking. Proceedings of the National Academy of Sciences 119(18), 2121153119 (2022)

[21] Agrawal, P., Singh, H., Srivastava, H.K., Singh, S., Kishore, G., Raghava, G.P.: Benchmarking of different molecular docking methods for protein-peptide docking. BMC bioinformatics 19(Suppl 13), 426 (2019)

[22] Porter, K.A., Xia, B., Beglov, D., Bohnuud, T., Alam, N., Schueler-Furman, O., Kozakov, D.: ClusPro PeptiDock: efficient global docking of peptide recognition motifs using FFT. Bioinformatics 33(20), 3299–3301 (2017)

[23] Blaszczyk, M., Ciemny, M.P., Kolinski, A., Kurcinski, M., Kmiecik, S.: Protein-peptide docking using CABS-dock and contact information. Briefings in bioinformatics 20(6), 2299–2305 (2019)

[24] Zhou, P., Jin, B., Li, H., Huang, S.-Y.: HPEPDOCK: a web server for blind peptide–protein docking based on a hierarchical algorithm. Nucleic acids research 46(W1), 443–450 (2018)

[25] Lee, H., Heo, L., Lee, M.S., Seok, C.: GalaxyPepDock: a protein–peptide docking tool based on interaction similarity and energy optimization. Nucleic acids research 43(W1), 431–435 (2015)

[26] Chen, T.S., Petrey, D., Garzon, J.I., Honig, B.: Predicting peptide-mediated interactions on a genome-wide scale. PLOS Computational Biology 11(5), 1–13 (2015). 10.1371/journal.pcbi.1004248

[27] Velez, C., Naravane, A., Robila, V.I., Saha, A., Murray, D., Honig, B.: Preppi – structure-based prediction of protein-protein interactomes and networks. Journal of Molecular Biology, 169735 (2026). 10.1016/j.jmb.2026.169735

[28] Jumper, J., Evans, R., Pritzel, A., Green, T., Figurnov, M., Ronneberger, O., Tunyasuvunakool, K., Bates, R., Žídek, A., Potapenko, A., et al.: Highly accurate protein structure prediction with AlphaFold. Nature 596(7873), 583–589 (2021)

[29] Lin, Z., Akin, H., Rao, R., Hie, B., Zhu, Z., Lu, W., Smetanin, N., Verkuil, R., Kabeli, O., Shmueli, Y., et al.: Evolutionary-scale prediction of atomic-level protein structure with a language model. Science 379(6637), 1123–1130 (2023)

[30] Baek, M., DiMaio, F., Anishchenko, I., Dauparas, J., Ovchinnikov, S., Lee, G.R., Wang, J., Cong, Q., Kinch, L.N., Schaeffer, R.D., et al.: Accurate prediction of protein structures and interactions using a three-track neural network. Science 373(6557), 871–876 (2021)

[31] Abramson, J., Adler, J., Dunger, J., Evans, R., Green, T., Pritzel, A., Ronneberger, O., Willmore, L., Ballard, A.J., Bambrick, J., et al.: Accurate structure prediction of biomolecular interactions with AlphaFold 3. Nature 630(8016), 493–500 (2024)

[32] Tsaban, T., Varga, J.K., Avraham, O., Ben-Aharon, Z., Khramushin, A., Schueler-Furman, O.: Harnessing protein folding neural networks for peptide-protein docking. Nature communications 13(1), 176 (2022)

[33] Motmaen, A., Dauparas, J., Baek, M., Abedi, M.H., Baker, D., Bradley, P.: Peptide-binding specificity prediction using fine-tuned protein structure prediction networks. Proceedings of the National Academy of Sciences 120(9), 2216697120 (2023)

[34] Hashemi, N., Hao, B., Ignatov, M., Paschalidis, I.C., Vakili, P., Vajda, S., Kozakov, D.: Improved prediction of MHC-peptide binding using protein language models. Frontiers in Bioinformatics 3, 1207380 (2023)

[35] Sledzieski, S., Singh, R., Cowen, L., Berger, B.: D-SCRIPT translates genome to phenome with sequence-based, structure-aware, genome-scale predictions of protein-protein interactions. Cell Systems 12(10), 969–982 (2021)

[36] Singh, R., Devkota, K., Sledzieski, S., Berger, B., Cowen, L.: Topsy-Turvy: integrating a global view into sequence-based PPI prediction. Bioinformatics 38(Supplement 1), 264–272 (2022)

[37] Sledzieski, S., Devkota, K., Singh, R., Cowen, L., Berger, B.: Tt3d: Leveraging precomputed protein 3d sequence models to predict protein–protein interactions. Bioinformatics 39(11), 663 (2023)

[38] Lei, Y., Li, S., Liu, Z., Wan, F., Tian, T., Li, S., Zhao, D., Zeng, J.: A deep-learning framework for multi-level peptide–protein interaction prediction. Nature communications 12(1), 5465 (2021)

[39] Cunningham, J.M., Koytiger, G., Sorger, P.K., AlQuraishi, M.: Biophysical prediction of protein– peptide interactions and signaling networks using machine learning. Nature methods 17(2), 175–183 (2020)

[40] Dens, C., Laukens, K., Bittremieux, W., Meysman, P.: The pitfalls of negative data bias for the T-cell epitope specificity challenge. Nature Machine Intelligence 5(10), 1060–1062 (2023)

[41] Grazioli, F., Mösch, A., Machart, P., Li, K., Alqassem, I., O’Donnell, T.J., Min, M.R.: On TCR binding predictors failing to generalize to unseen peptides. Frontiers in immunology 13, 1014256 (2022)

[42] Elnaggar, A., Heinzinger, M., Dallago, C., Rehawi, G., Wang, Y., Jones, L., Gibbs, T., Feher, T., Angerer, C., Steinegger, M., et al.: Prottrans: Toward understanding the language of life through self-supervised learning. IEEE transactions on pattern analysis and machine intelligence 44(10), 7112–7127 (2021)

[43] Lin, Z., Akin, H., Rao, R., Hie, B., Zhu, Z., Lu, W., Smetanin, N., Verkuil, R., Kabeli, O., Shmueli, Y., et al.: Evolutionary-scale prediction of atomic-level protein structure with a language model. Science 379(6637), 1123–1130 (2023)

[44] Brandes, N., Ofer, D., Peleg, Y., Rappoport, N., Linial, M.: ProteinBERT: a universal deep-learning model of protein sequence and function. Bioinformatics 38(8), 2102–2110 (2022)

[45] Elnaggar, A., Essam, H., Salah-Eldin, W., Moustafa, W., Elkerdawy, M., Rochereau, C., Rost, B.: Ankh: Optimized protein language model unlocks general-purpose modelling. arXiv preprint arXiv:2301.06568 (2023)

[46] Majila, K., Ullanat, V., Viswanath, S.: Disobind: A sequence-based, partner-dependent contact map and interface residue predictor for intrinsically disordered regions. Cell Systems 17(1) (2026)

[47] Wang, M., Patsenker, J., Li, H., Kluger, Y., Kleinstein, S.H.: Language model-based B cell receptor sequence embeddings can effectively encode receptor specificity. Nucleic acids research 52(2), 548–557 (2024)

[48] Wang, Y., Lv, H., Teo, Q.W., Lei, R., Gopal, A.B., Ouyang, W.O., Yeung, Y.-H., Tan, T.J., Choi, D., Shen, I.R., et al.: An explainable language model for antibody specificity prediction using curated influenza hemagglutinin antibodies. Immunity 57(10), 2453–2465 (2024)

[49] Schmirler, R., Heinzinger, M., Rost, B.: Fine-tuning protein language models boosts predictions across diverse tasks. Nature Communications 15(1), 7407 (2024)

[50] Sledzieski, S., Kshirsagar, M., Baek, M., Dodhia, R., Ferres, J.L., Berger, B.: Democratizing protein language models with parameter-efficient fine-tuning. Proceedings of the National Academy of Sciences 121(26), 2405840121 (2024). 10.1073/pnas.2405840121

[51] Zhou, H., Yin, M., Wu, W., Li, M., Fu, K., Chen, J., Wu, J., Wang, Z.: ProtCLIP: Function-informed protein multi-modal learning. In: Proceedings of the AAAI Conference on Artificial Intelligence, vol. 39, pp. 22937–22945 (2025)

[52] Im, C., Zhao, R., Boyd, S.D., Kundaje, A.: Sequence-based TCR-peptide representations using cross-epitope contrastive fine-tuning of protein language models. In: International Conference on Research in Computational Molecular Biology, pp. 34–48 (2025). Springer

[53] Bhat, S., Palepu, K., Hong, L., Mao, J., Ye, T., Iyer, R., Zhao, L., Chen, T., Vincoff, S., Watson, R., et al.: De novo design of peptide binders to conformationally diverse targets with contrastive language modeling. Science Advances 11(4), 8638 (2025)

[54] Cohen, T., Schneidman-Duhovny, D.: Epitope-specific antibody design using diffusion models on the latent space of ESM embeddings. In: ICLR 2024 Workshop on Generative and Experimental Perspectives for Biomolecular Design (2023)

[55] Jia, Y., Gao, B., Tan, J., Zheng, J., Hong, X., Zhu, W., Tan, H., Xiao, Y., Tan, L., Cai, H., Huang, Y., Deng, Z., Wu, X., Jin, Y., Yuan, Y., Tian, J., He, W., Ma, W., Zhang, Y., Liu, L., Yan, C., Zhang, W., Lan, Y.: Deep contrastive learning enables genome-wide virtual screening. Science 391(6781), 9530 (2026). 10.1126/science.ads9530

[56] Im, C., Zhao, R., Boyd, S.D., Kundaje, A.: Sequence-based TCR-peptide representations using cross-epitope contrastive fine-tuning of protein language models. In: International Conference on Research in Computational Molecular Biology, pp. 34–48 (2025). Springer

[57] Singh, R., Sledzieski, S., Bryson, B., Cowen, L., Berger, B.: Contrastive learning in protein language space predicts interactions between drugs and protein targets. Proceedings of the National Academy of Sciences 120(24), 2220778120 (2023)

[58] Jha, K., Shonai, D., Parekh, A., Uezu, A., Fujiyama, T., Yamamoto, H., Parameswaran, P., Yanagisawa, M., Singh, R., Soderling, S.H.: Deep learning-coupled proximity proteomics to deconvolve kinase signaling in vivo. bioRxiv (2025). 10.1101/2025.04.27.650849

[59] Cornman, A., Tranzillo, M., Zulaybar, N.G., Bouzit, I., Hwang, Y.: Linear-time prediction of proteome-scale microbial protein interactions. bioRxiv (2026). 10.64898/2026.03.01.708874

[60] Dapkūnas, J., Timinskas, A., Olechnovič, K., Margelevičius, M., Dičiūnas, R., Venclovas, Č.: The PPI3D web server for searching, analyzing and modeling protein–protein interactions in the context of 3D structures. Bioinformatics 33(6), 935–937 (2017)

[61] Kovtun, D., Akdel, M., Goncearenco, A., Zhou, G., Holt, G., Baugher, D., Lin, D., Adeshina, Y., Castiglione, T., Wang, X., Marquet, C., McPartlon, M., Geffner, T., Rossi, E., Corso, G., Stärk, H., Carpenter, Z., Kucukbenli, E., Bronstein, M., Naef, L.: PINDER: The protein interaction dataset and evaluation resource. bioRxiv (2024). 10.1101/2024.07.17.603980

[62] London, N., Raveh, B., Movshovitz-Attias, D., Schueler-Furman, O.: Can self-inhibitory peptides be derived from the interfaces of globular protein–protein interactions? Proteins: Structure, Function, and Bioinformatics 78(15), 3140–3149 (2010)

[63] Vanhee, P., Stricher, F., Baeten, L., Verschueren, E., Lenaerts, T., Serrano, L., Rousseau, F., Schymkowitz, J.: Protein-peptide interactions adopt the same structural motifs as monomeric protein folds. Structure 17(8), 1128–1136 (2009). 10.1016/j.str.2009.06.013

[64] Radford, A., Kim, J.W., Hallacy, C., Ramesh, A., Goh, G., Agarwal, S., Sastry, G., Askell, A., Mishkin, P., Clark, J., et al.: Learning transferable visual models from natural language supervision. In: International Conference on Machine Learning, PMLR, pp. 8748–8763 (2021)

[65] ESM Team: ESM Cambrian: Revealing the mysteries of proteins with unsupervised learning. EvolutionaryScale Website (2024). https://evolutionaryscale.ai/blog/esm-cambrian

[66] Olechnovič, K., Venclovas, Č.: Voronota: a fast and reliable tool for computing the vertices of the Voronoi diagram of atomic balls. Journal of computational chemistry 35(8), 672–681 (2014)

[67] Lavi, A., Ngan, C.H., Movshovitz-Attias, D., Bohnuud, T., Yueh, C., Beglov, D., Schueler-Furman, O., Kozakov, D.: Detection of peptide-binding sites on protein surfaces: The first step toward the modeling and targeting of peptide-mediated interactions. Proteins: Structure, Function, and Bioinformatics 81(12), 2096–2105 (2013)

[68] Abdin, O., Nim, S., Wen, H., Kim, P.M.: PepNN: a deep attention model for the identification of peptide binding sites. Communications biology 5(1), 503 (2022)

[69] Hu, E.J., Shen, Y., Wallis, P., Allen-Zhu, Z., Li, Y., Wang, S., Wang, L., Chen, W., et al.: Lora: Low-rank adaptation of large language models. ICLR 1(2), 3 (2022)

[70] Varga, J.K., Ovchinnikov, S., Schueler-Furman, O.: actifpTM: a refined confidence metric of AlphaFold2 predictions involving flexible regions. Bioinformatics 41(3), 107 (2025)

[71] Dunbrack, R.L.: Res ipSAE loquunt: What’s wrong with AlphaFold’s ipTM score and how to fix it. bioRxiv (2025). 10.1101/2025.02.10.637595

[72] Kliche, J., Garvanska, D.H., Simonetti, L., Badgujar, D., Dobritzsch, D., Nilsson, J., Davey, N.E., Ivarsson, Y.: Large-scale phosphomimetic screening identifies phospho-modulated motif-based protein interactions. Molecular Systems Biology 19(7), 202211164 (2023). 10.15252/msb.202211164

[73] Mihalič, F., Simonetti, L., Giudice, G., Sander, M.R., Lindqvist, R., Peters, M.B.A., Benz, C., Kassa, E., Badgujar, D., Inturi, R., et al.: Large-scale phage-based screening reveals extensive pan-viral mimicry of host short linear motifs. Nature Communications 14(1), 2409 (2023)

[74] Benz, C., Ali, M., Krystkowiak, I., Simonetti, L., Sayadi, A., Mihalic, F., Kliche, J., Andersson, E., Jemth, P., Davey, N.E., et al.: Proteome-scale mapping of binding sites in the unstructured regions of the human proteome. Molecular Systems Biology 18(1), 10584 (2022)

[75] Kosugi, S., Hasebe, M., Tomita, M., Yanagawa, H.: Nuclear export signal consensus sequences defined using a localization-based yeast selection system. Traffic 9(12), 2053–2062 (2008)

[76] Evans, R., O’Neill, M., Pritzel, A., Antropova, N., Senior, A., Green, T., Žídek, A., Bates, R., Blackwell, S., Yim, J., Ronneberger, O., Bodenstein, S., Zielinski, M., Bridgland, A., Potapenko, A., Cowie, A., Tunyasuvunakool, K., Jain, R., Clancy, E., Kohli, P., Jumper, J., Hassabis, D.: Protein complex prediction with AlphaFold-Multimer (2021). 10.1101/2021.10.04.463034

[77] Kehlenbach, R.H., Chook, Y.M.: The nuclear export receptor CRM1/XPO1 and its diverse cargoes. Trends in Biochemical Sciences 50(12), 1131–1144 (2025). 10.1016/j.tibs.2025.09.003

[78] Sendino, M., Omaetxebarria, M.J., Prieto, G., Rodriguez, J.A.: Using a simple cellular assay to map NES motifs in cancer-related proteins, gain insight into CRM1-mediated NES export, and search for NES-harboring micropeptides. International journal of molecular sciences 21(17), 6341 (2020)

[79] UniProt Proteome UP000005640 (Homo sapiens). https://www.uniprot.org/proteomes/UP000005640. Accessed: 2025-07-14

[80] Kırlı, K., Karaca, S., Dehne, H.J., Samwer, M., Pan, K.T., Lenz, C., Urlaub, H., Görlich, D.: A deep proteomics perspective on CRM1-mediated nuclear export and nucleocytoplasmic partitioning. elife 4, 11466 (2015)

[81] Landrum, M.J., Chitipiralla, S., Brown, G.R., Chen, C., Gu, B., Hart, J., Hoffman, D., Jang, W., Kaur, K., Liu, C., et al.: ClinVar: improvements to accessing data. Nucleic acids research 48(D1), 835–844 (2020)

[82] Brunello, F.G., Erra, L., Nicola, J., Martí, M.A.: Integrating AlphaFold2 models and clinical data to improve the assessment of Short Linear Motifs (SLiMs) and their variants’ pathogenicity. PLOS Computational Biology 21(8), 1012829 (2025)

[83] Xiong, D., Qiu, Y., Zhao, J., Zhou, Y., Lee, D., Gupta, S., Torres, M., Lu, W., Liang, S., Kang, J.J., et al.: A structurally informed human protein–protein interactome reveals proteome-wide perturbations caused by disease mutations. Nature Biotechnology, 1–15 (2024)

[84] Livesey, B.J., Marsh, J.A.: The properties of human disease mutations at protein interfaces. PLOS Computational Biology 18(2), 1009858 (2022)

[85] Berman, H.M., Westbrook, J., Feng, Z., Gilliland, G., Bhat, T.N., Weissig, H., Shindyalov, I.N., Bourne, P.E.: The Protein Data Bank. Nucleic Acids Research 28(1), 235–242 (2000). 10.1093/nar/28.1.235

[86] Wells, J., Hawkins-Hooker, A., Bordin, N., Sillitoe, I., Paige, B., Orengo, C.: Chainsaw: protein domain segmentation with fully convolutional neural networks. Bioinformatics 40(5), 296 (2024)

[87] Mirdita, M., Schütze, K., Moriwaki, Y., Heo, L., Ovchinnikov, S., Steinegger, M.: ColabFold: making protein folding accessible to all. Nature methods 19(6), 679–682 (2022)

