## Supplementary Info for "CliPepPI: Scalable prediction of domain-peptide specificity using contrastive learning"

---

#### Contents

**Supplementary Figures** **2**

**Supplementary Tables** **6**

### Supplementary Figures

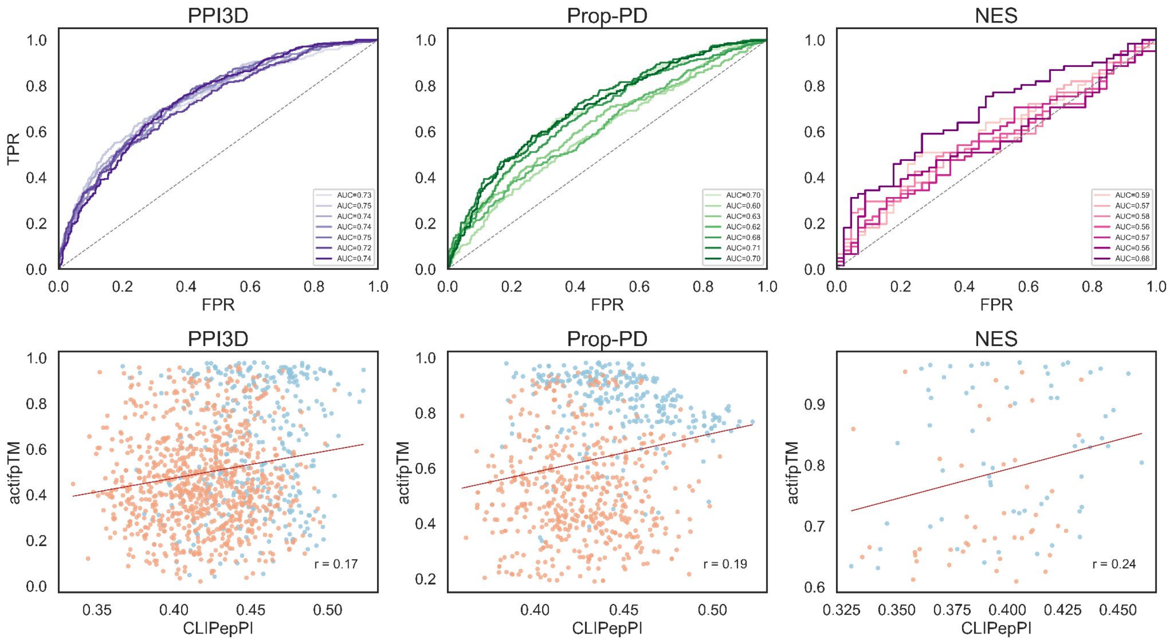


Figure S1: **CLIPepPI-average-loss performance across three benchmark datasets. A.** Receiver-operating-characteristic (ROC) curves for the seven cross-validated models on the PPI3D, ProP-PD, and NES test sets (left to right). Each shaded curve corresponds to a single cross-validation model; the dashed diagonal denotes random classification. Legends list the per-model area-under-the-curve (AUC) values. Note the better performance on the PPI3D test set, but worse performance on the Prop-PD and NES test sets, compared to our weighted-loss models (Fig. 2C). **B.** Scatter plots comparing CLIPepPI-average-loss scores (x-axis) with AlphaFold inter-chain confidence scores (*actifpTM*; y-axis) for the same datasets. Blue dots represent positive (binding) pairs and orange dots negative pairs.

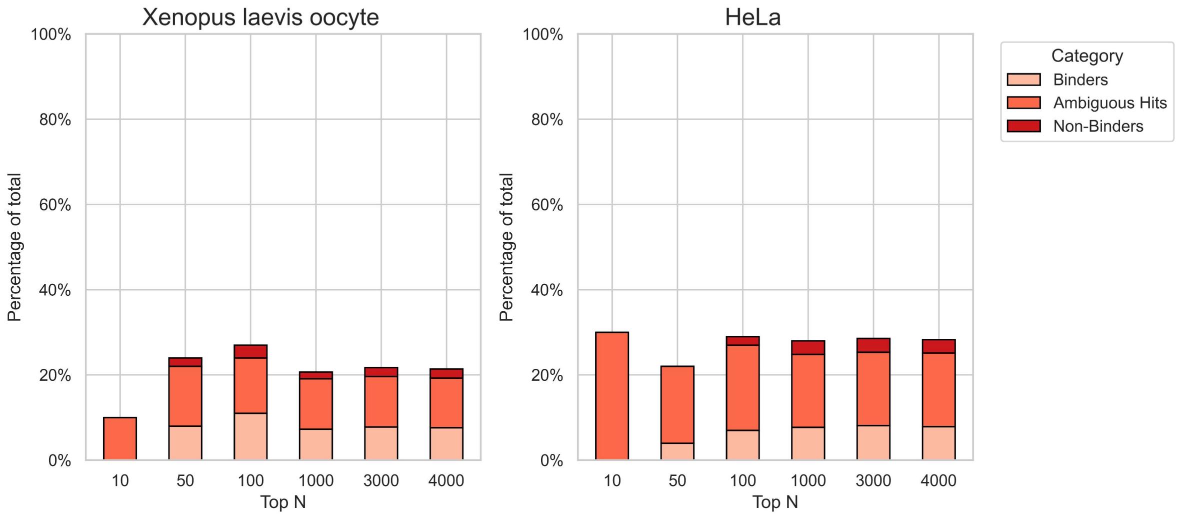


Figure S2: **Proteome-wide scan using the CLIPepPI-average-loss for NES motif containing proteins compared against experimental proteomics annotations *HeLa* cells and *Xenopus laevis* oocytes.** For increasing Top-$N$ cutoffs, stacked bars report the percentage of total hits among the highest-scoring candidates, partitioned by annotation categories from proteomics experiments (see legend). Compare to Fig. 3B in the main text.

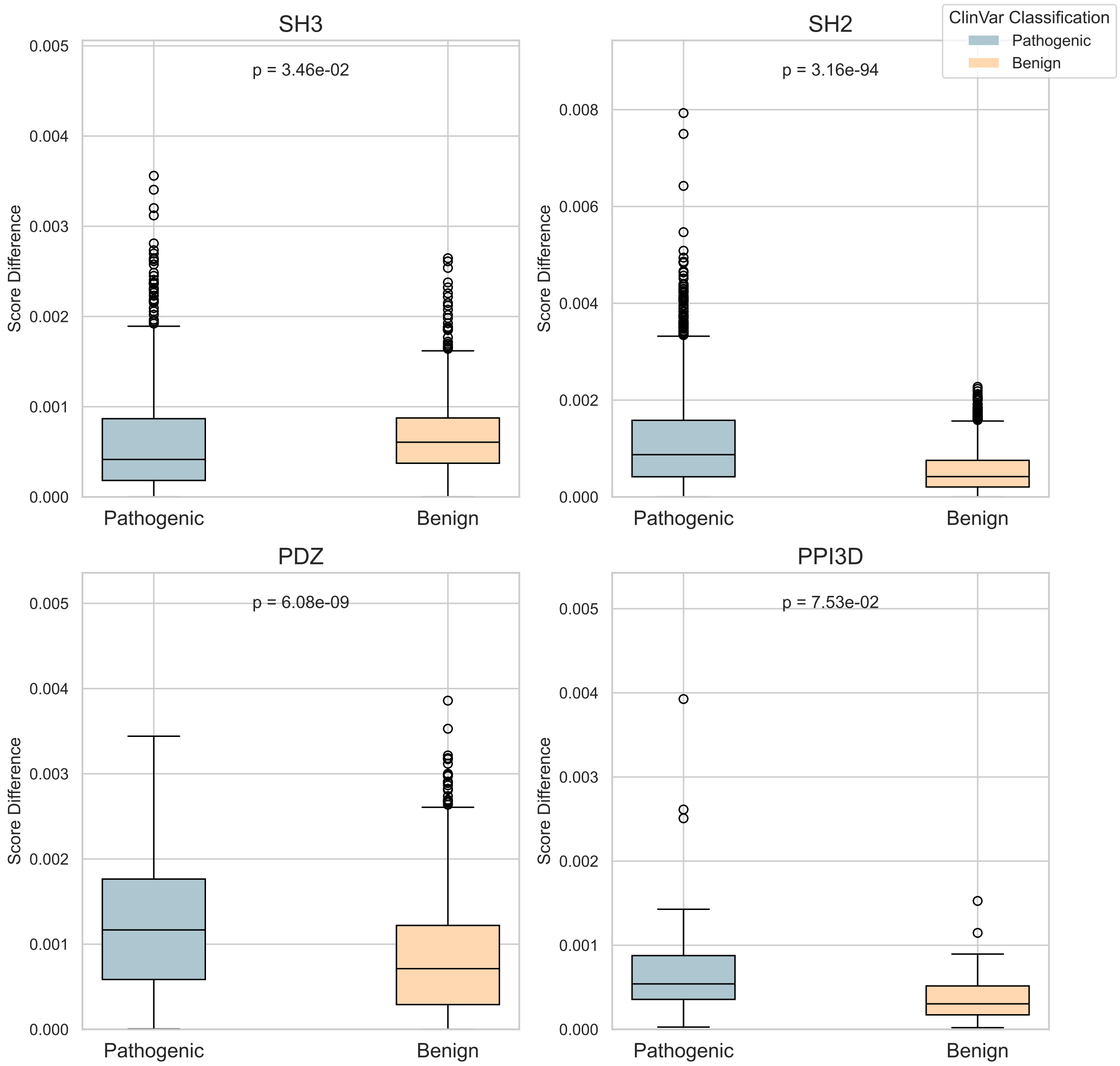


Figure S3: **Analysis of ClinVar variants for CLIPepPI-average-loss model.** Analysis of effects of ClinVar variants. Distributions of score difference (wildtype vs. mutant) for three domain families (SH3, WW, PDZ) and general domains (PPI3D). Boxplots show the median, interquartile range; points are outliers. T-test *p*-values are annotated above each panel, indicating a significant separation between the score differences of pathogenic and benign variants. Compare to Fig. 4 in the main text

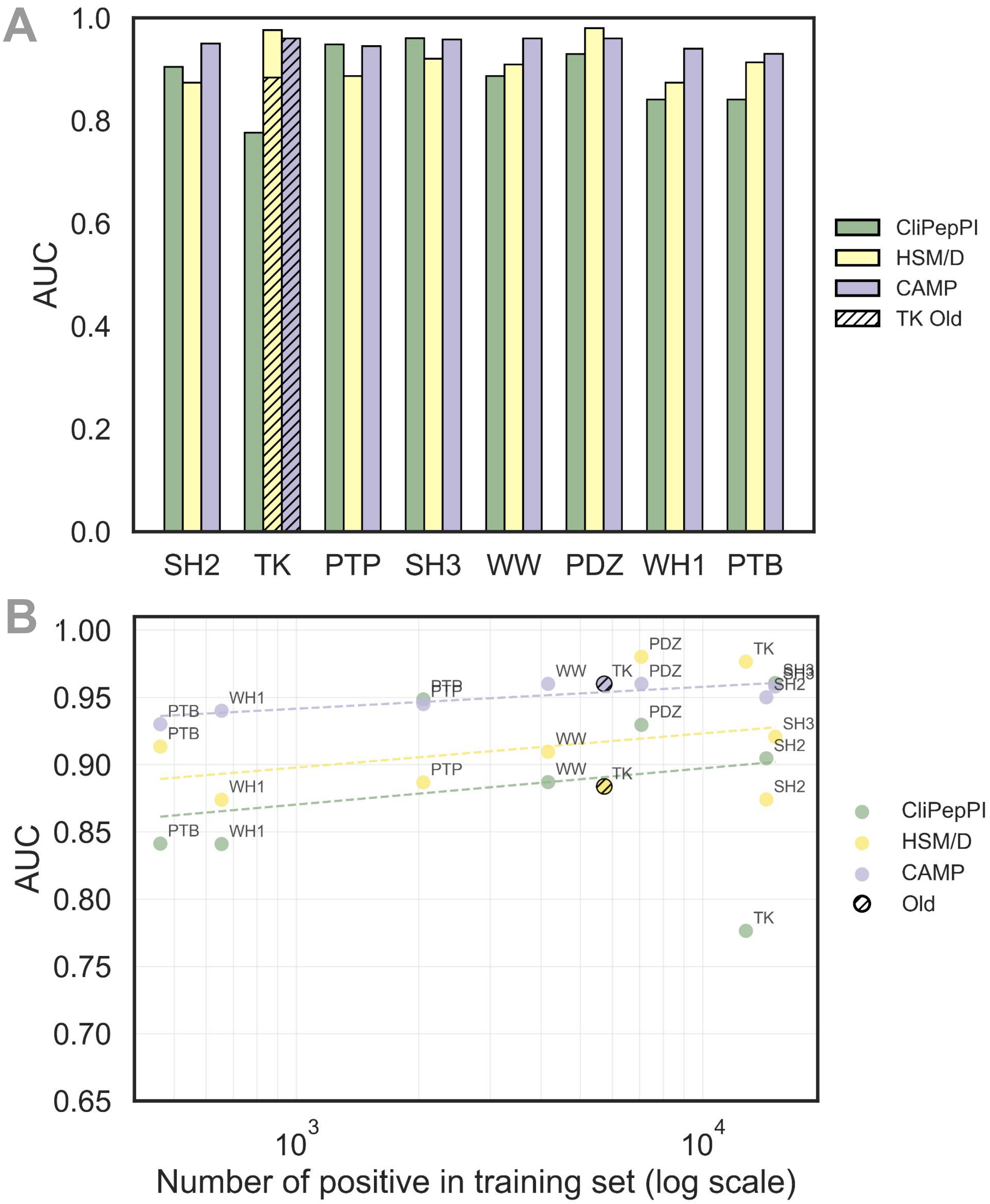


Figure S4: **Comparison of domain-peptide binding prediction models across domain families.** **A.** AUC scores for several benchmark models: **CLIPepPI**, **CAMP**, and **HSM/D**, evaluated on diverse domain families including SH2, SH3, PDZ, WW, PTB, PTP, TK, and WH1. A revised TK dataset was released by Cunningham *et al.* in 2023. Since CAMP was trained and evaluated on the older dataset, we report HSM/D performance on both datasets and denote old results with hatched lines. Each group of bars represents the performance of the different models on a specific domain type, highlighting the strengths and weaknesses of each method. CLIPepPI outperforms on part of the families with high results on SH3 and PTP. **B.** Model performance versus positive train data size across domain families. Hatched dots denote old TK dataset size.

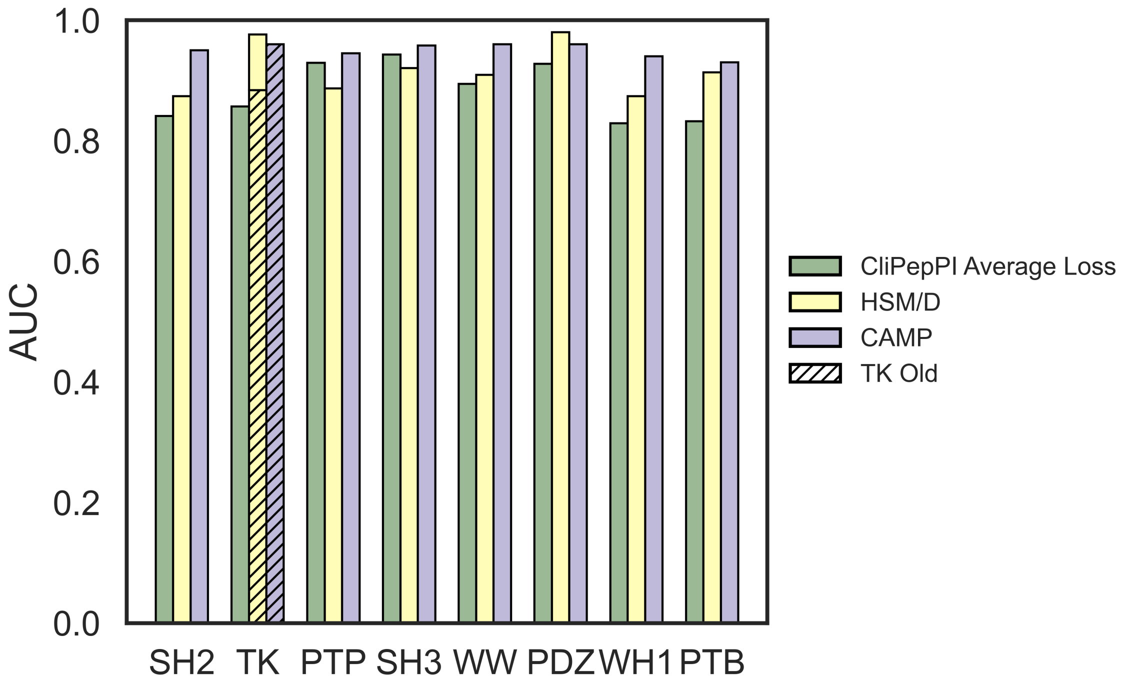


Figure S5: **Comparison of domain-peptide binding prediction models across domain families using CLIPepPI-average-loss model.** AUC scores for several benchmark models: **CLIPepPI**, **CAMP**, and **HSM/D**, evaluated on diverse domain families including SH2, SH3, PDZ, WW, PTB, PTP, TK, and WH1. A revised TK dataset was released by Cunningham *et al.* in 2023. Since CAMP was trained and evaluated on the older dataset, we report HSM/D performance on both datasets and denote old results with hatched lines. Each group of bars represents the performance of the different models on a specific domain type, highlighting the strengths and weaknesses of each method.

### Supplementary Tables

| Hyperparameter | |
|---|---|
| # epochs | 200 |
| batch size | 16 ~ |
| learning rate | 0.0001 |
| optimizer | AdamW |
| dropout rate | 0.15 |
| LoRA alpha | 32 |
| LoRA r | 8 |

Table 1: Training hyper-parameters
